# Structural divergence in protein evolution of a photoreceptor family undetected by AlphaFold is observed by high sensitivity FT-IR spectroscopy

**DOI:** 10.64898/2026.08.04.742864

**Authors:** Rosalie L. Dohmen, Gunnar Hoogerwerf, Aihua Xie, Wouter D. Hoff

## Abstract

A universal mechanism in molecular evolution is functional and structural divergence of members of a protein family. The ability of AlphaFold to predict atomic-resolution protein structures promises to accelerate insights into this process. We study the interplay of changes in sequence, structure, and function in photoactive yellow protein (PYP), a family of bacterial blue light photoreceptors. *Halorhodospira halophila* contains two PYP homologs that diverged to 60% sequence identity, differ 100-fold in the lifetime (τ_pB_) of their pB signaling intermediate, and display altered peak wavelengths (λ_max_) for color sensing. We resurrected ancestral PYPs and determined these properties along the resulting recapitulating evolutionary divergence. The resurrected ancestral PYP is functionally similar to PYP1, indicating divergence on the path to PYP2. AlphaFold predictions for PYP2 and these ancestral proteins revealed the absence of structural changes compared to the crystal structure of PYP1. To experimentally validate these predictions, we optimized second-derivative Fourier transform infrared (FTIR) spectroscopy. The FTIR spectra of PYP1 and 2 and their resurrected ancestral proteins demonstrated clear differences in their secondary structure. These results demonstrate an important limitation of AlphaFold and show how ancestral sequence reconstruction combined with spectroscopic approaches yields insights into divergence in a protein family.

## Introduction

Functional divergence of proteins is a major contributor to the process of molecular evolution ^1–5^. This process results in the very large numbers of homologs now known for many protein families ^6^. Here we study such divergence in the Photoactive Yellow Protein (PYP) family of bacterial photoreceptors ^7–9^. PYP was first discovered in the photosynthetic bacterium *Halorhodospira halophila* as a small (125 residues, 14 kDa), water-soluble, cytoplasmic photoreceptor ^10,11^. PYP has been reported to serve as the photoreceptor that triggers a negative response to blue light in this organism ^12^. PYP is widely regarded as a model system for investigating functional protein dynamics and signaling within the PAS domain superfamily ^13–16^. It contains a *p*-coumaric acid (*p*CA) chromophore that is covalently bound to cysteine 69 via thioester bond ^17–20^ and its three-dimensional structure has been reported at high resolution ^21–23^. In the ground state (pG), PYP1 exhibits an absorbance maximum (λ_max_) at 447 nm. Upon photoexcitation, the central C=C bond of the *p*CA undergoes *trans* to *cis* photoisomerization ^24^, resulting in the formation of a blue-shift photocycle intermediate (pB) with the λ_max_ near 355 nm. The pB intermediate is thought to be the signaling state of PYP, and it thermally returns to the initial pG state with a lifetime τ_pB_ of approximately 0.5 seconds ^11,25^. The exact value of τ_pB_ varies depending on the temperature, pH, and buffer conditions used^11,26,27^. The light-triggered nature of PYPs provides superb experimental accessibility to the conformational changes associated with PYP function, thereby facilitating the exploration of structure-function relationships ^28,29^. While the PYP from *H. halophila* remains the best-studied PYP, 12 additional PYP homologs have been studied, revealing considerable divergence in their functional properties, particularly their absorbance maxima and photocycle kinetics.^7–9,28,30–37,37–39^. While the interactions between the *p*CA chromophore and residues in the active site of PYP that govern the position of that govern the position of λ_ma_ ^40–47^, the mechanisms that determine the value of τ_pB_ of remain poorly understood.

20 years after the discovery of PYP in *H. halophila*, biochemical experiments based on genome sequence information ^48^ revealed that this organism possesses two distinct PYPs ^49^, which are referred to here as PYP1 and PYP2. While PYP2 remains much less studied than PYP1, it displays considerable divergence in its amino acid sequence and also exhibits distinct functional characteristics. PYP1 and PYP2 share sequence identity of 60% and similarity of 74%, differing at 50 positions. With respect to biochemical properties, PYP2 shows a λ_max_ around 442 nm, and a thermal decay of its pB intermediate to the pG state of approximately 1 minute, some 100-fold slower than in PYP1 (Figure 1). The physiological function of PYP2 remains unknown, but it’s much slower overall photocycle rate suggests the possibility that it photo-regulates slower responses in the cell ^34,50^. PYP1 and PYP2 from *H. halophila* provide an opportunity to study the molecular mechanism of how molecular evolution results in the divergence of functional properties within a protein family, both with respect to active site interactions (λ_max_) and functional kinetics (τ_pB_).

**Figure 1.**
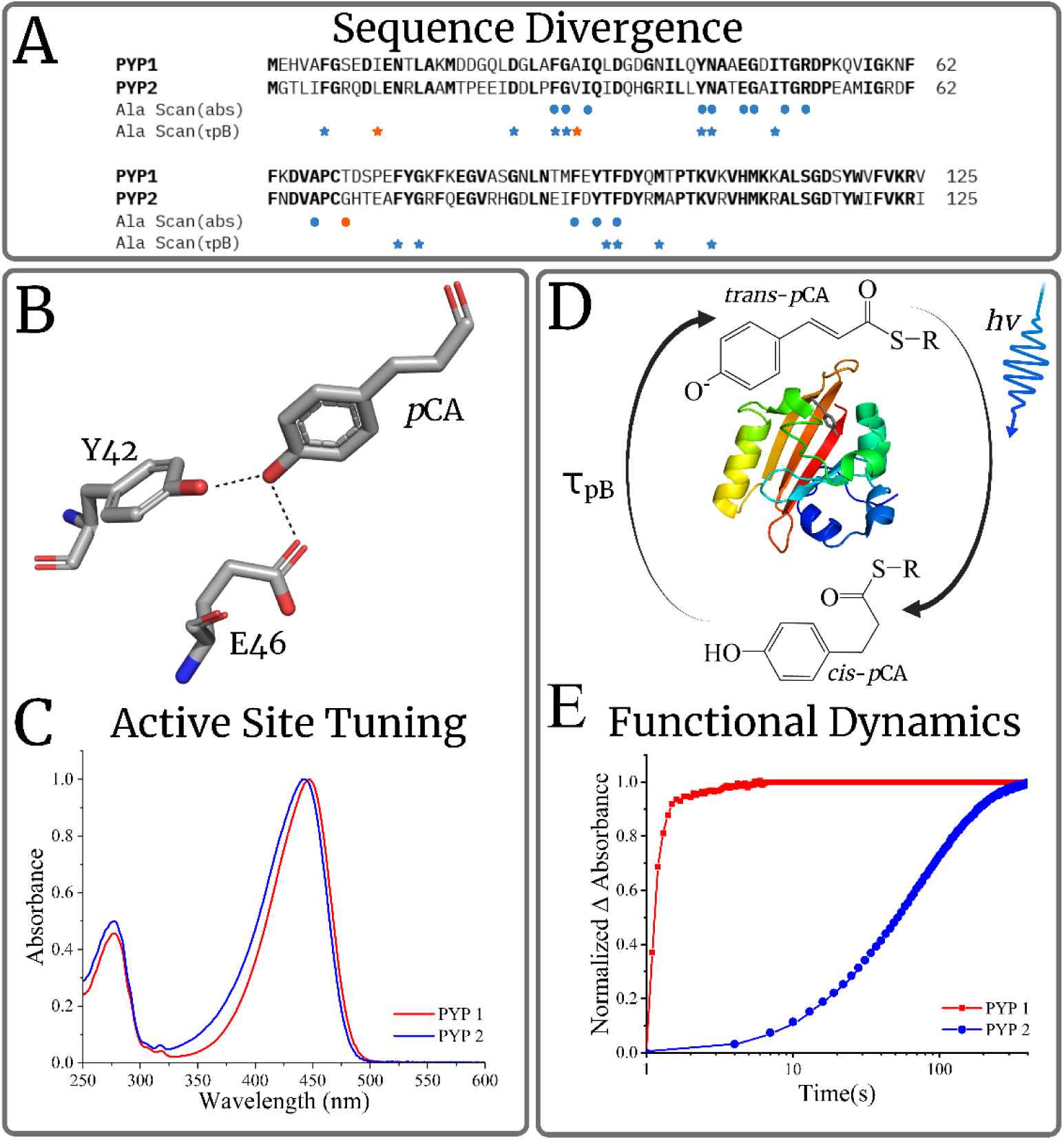
Evolutionary divergence of the functional properties of PYP1 and PYP2 from *Halorhodospira halophila*. The two PYP homologs from *H. halophila* exhibit substantial sequence diversity (A) and functional divergence (B-D). (A) Sequence alignment of PYP1 and PYP2. Residues identified by Ala scan mutagenesis data of PYP1 as affecting λ_max_ by at least 3 nanometers (circles) or τ_pB_ (stars) by at least a factor 10 are indicated. Conserved residues are indicated in bold. (B) Structure of the photo-active site of PYP illustrating hydrogen bonds between the *p*CA chromophore and active site residues Tyr42 and Glu46 that are involved in functional tuning. (C) Illustration of spectral tuning of the *p*CA absorbance peak in PYP. (D) Simplified scheme of the PYP photocycle depicting the photoconversion of the initial pG dark state (λ_max_ 447 nm) to the long-lived pB photocycle intermediate (λ_max_ 355 nm) after light-induced *trans* to *cis* isomerization and protonation of the *p*CA chromophore. The pB intermediate spontaneously decays back to pG with a time constant τ_pB._ (E) Time-resolved UV/Vis absorbance measurements at 446 nm during pB decay allow quantification of τ_pB_.

The phenomenon that proteins can shift the λ_max_ of a chromophore by altering protein-chromophore interactions is referred to as spectral tuning. Such spectral tuning has attracted considerable attention because it is at the basis of human color vision ^51–53^ and also has been investigated in PYP1 ^43–46,54,55^. As a result, a number of point mutations affecting the position of the λ_max_ of PYP1 have been identified ^28,29,46^. The strength of the active-site hydrogen bond between the *p*CA chromophore and the protonated side chain of Tyr42 ^40,56^ and Glu46 ^43,45^ has been identified as being directly involved in tuning the position of the visible absorbance peak of PYP ^43^. Regarding τ_pB_, multiple point mutations have been identified ^28,29^ that alter the rate of pB decay over a 10,000-fold range, but the mechanism for these effects has not yet been uncovered. Figure 1 provides a summary of this information on structure-function relationships in PYP.

Here we use ancestral sequence reconstruction (ASR) to study functional divergence in the PYP family of photoreceptors. The use of experimental studies on resurrected ancestral proteins with sequences that were computed using bioinformatics techniques has come to the forefront as a powerful approach to study such questions that combine evolutionary and mechanistic aspects^57–60^. These computationally derived ancestral proteins are designed to recapitulate the evolutionary process of protein diversification, and are predicted to be proteins that were functional in ancestral organisms. We used Alphafold2, a tool that has dramatically come to the forefront of protein science ^61,62^, to predict the structures of the ancestral PYPs resurrected here. To validate these structural predictions, we used FTIR spectroscopy, a powerful spectroscopic approach to study protein structural features ^16,63^. Surprisingly, we find that substantial changes in protein structure during the reconstructed evolutionary pathway that we determine experimentally using FTIR spectroscopy were entirely missed by the computational predictions. Together, these results provide novel insights into the mechanism of protein evolution and indicate that important structural changes during the evolution of proteins are not adequately captured by Alphafold2 structure predictions.

## Materials and Methods

### Bioinformatics analyses

The sequence of PYP1 from *H. halophila* (PDB: 1NWZ) was used as the query for a NCBI PSI-BLAST search against the non-redundant database to retrieve candidate PYP homologs. A high-quality multiple sequence alignment (MSA) was generated using MAFFT version 7.490 ^64^ and refined using CIAlign version 1.0.17 ^65^. The resulting alignment was visually inspected to confirm the presence of residues known to be functionally important for PYP, and sequences lacking Cys69 were removed. This MSA was used to generate a maximum likelihood phylogenetic tree using IQ-TREE 1.16.12 ^66,67^ with PAS domain protein PDB: 3FG8 from *Rhodococcus jostii* RHA1 as an outgroup and also to obtain predicted ancestral sequences using an empirical Bayesian method. For selected PYPs, ColabFold ^62^ was used to obtain AlphaFold2 structure predictions. The RMSD was computed using the Dali Server^68^ and the RMSF value was computed using ChimeraX 1.9 ^69^.

### Molecular biology and protein purification of PYPs

The genes encoding ancestral and extant PYPs were synthesized by Genscript, cloned into pET28b+ between the NdeI and BamHI restriction sites, transformed into NEB® 5-alpha Competent *E. coli* (High Efficiency) cells, and checked for DNA sequence. Subsequent PCR amplification and Gibson assembly were used to remove the N-terminal histidine tag, and the resulting constructs were transformed into *E. coli* strain BL21 (DE3). All PYPs were overproduced and purified as previously described in ^70^. After growth and induction with IPTG, cells were pelleted and extracted with 8 M urea. The cell free extract containing apoPYP was diluted 2-fold using 10 mM Tris-HCl buffer at pH 7.5 and reconstituted with the *p*-coumaric acid anhydride as described previously ^71^. After dialysis against 10 mM Tris-HCl buffer at pH 7.5, the protein was purified by three Sepharose anion exchange chromatography steps until the purity index (ratio of the absorption at 446 nm and 278 nm) was <0.45.

### UV/Vis absorbance spectroscopy

All UV/Vis absorbance measurements were performed using a HP-8453 (Hewlett-Packard) diode array spectrophotometer. Absorbance spectra were measured in standard mode. Spectra and photocycle kinetics during pB decay were measured in kinetic mode after illumination of the PYP sample using a 150-watt halogen quartz light source (Cuda) equipped with a light guide. The extinction coefficient of PYP was determined by measuring the absorbance spectrum before and after denaturation with 2% SDS as described previously ^40^.

### FTIR spectroscopy

Following recently reported methods ^70^. PYP samples were concentrated in 10 mM potassium phosphate buffer pH 7.5 to a protein concentration of 5 mM using 0.5ml Amicon centrifugal filters (Millipore Sigma) with a molecular weight cutoff of 10,000. Samples were then freeze dried overnight and rehydrated with half of the initial volume using pure D_2_O (to achieve a PYP concentration of 10 mM) just before FTIR measurements. FTIR spectra were collected at 2 cm^-1^ resolution from 950 to 4000 cm^-1^ using a Bruker Vertex 80V FTIR spectrometer with a KBr beamsplitter and a liquid nitrogen cooled photovoltaic mercury cadmium telluride (MCT) infrared detector. An AquaSpec (Bruker) FTIR sample cell with fixed sample thickness (∼7.0 μm) was used in all data collection, using purging with nitrogen gas of the sample cell and evacuation of the remaining optical path. FTIR data processing was performed in Bruker OPUS software by averaging spectra, performing a baseline shift (using a single offset value), then normalizing the amplitude of the resulting spectrum, and finally calculating its second derivative using the Savitsky-Golay algorithm over 17 data points.

## Results and Discussion

### Ancestral sequence reconstruction of PYP

The information needed for computing the amino acid sequences of ancestral proteins is derived from the sequence information of extant homologs. We therefore used a manually curated high-quality set of 984 extant PYP homologs that was recently reported ^7^. Each of these sequences was manually inspected for the presence of functionally important and highly conserved residues ^7–9,28^, and was confirmed to contain the functionally essential residue Cys69 ^20,29^. Using this high-quality multiple sequence alignment (MSA), we reconstructed a full set of ancestral proteins describing the diversification of the PYP family. From this series of reconstructed ancestral PYPs, we extracted the last common ancestor (LCA) of PYP1 and PYP2. This LCA differs at 34 positions from PYP1 and at 31 positions from PYP2 (see Supplemental Figure 1). The set of PYP homologs we recovered allowed us to compute a total of 11 ancestors in the transition from the LCA to PYP1 and one ancestor on the path from the LCA to PYP2. We selected this single intermediate ancestor (referred to as Node 2A) in between the LCA and PYP2 for experiment resurrection together with two intermediates (Nodes 1A and 1B) on the pathway from the LCA to PYP1. The latter two ancestors were chosen to have approximately equidistant steps (between 10 and 16 substitutions) in sequence space (Figure 2). Thus, the computed phylogenetic tree of PYPs and the computed ancestral proteins that yielded the diversity of extant PYPs provide a stepwise path through sequence space that connects PYP1 and PYP2 via series of ancestors that are all predicted to be functional PYPs (Figure 2, bottom panel). The ancestral nodes exhibit percentage identities to PYP1 and PYP2 that range from 58.4% to 92% (Table 1).

**Figure 2.**
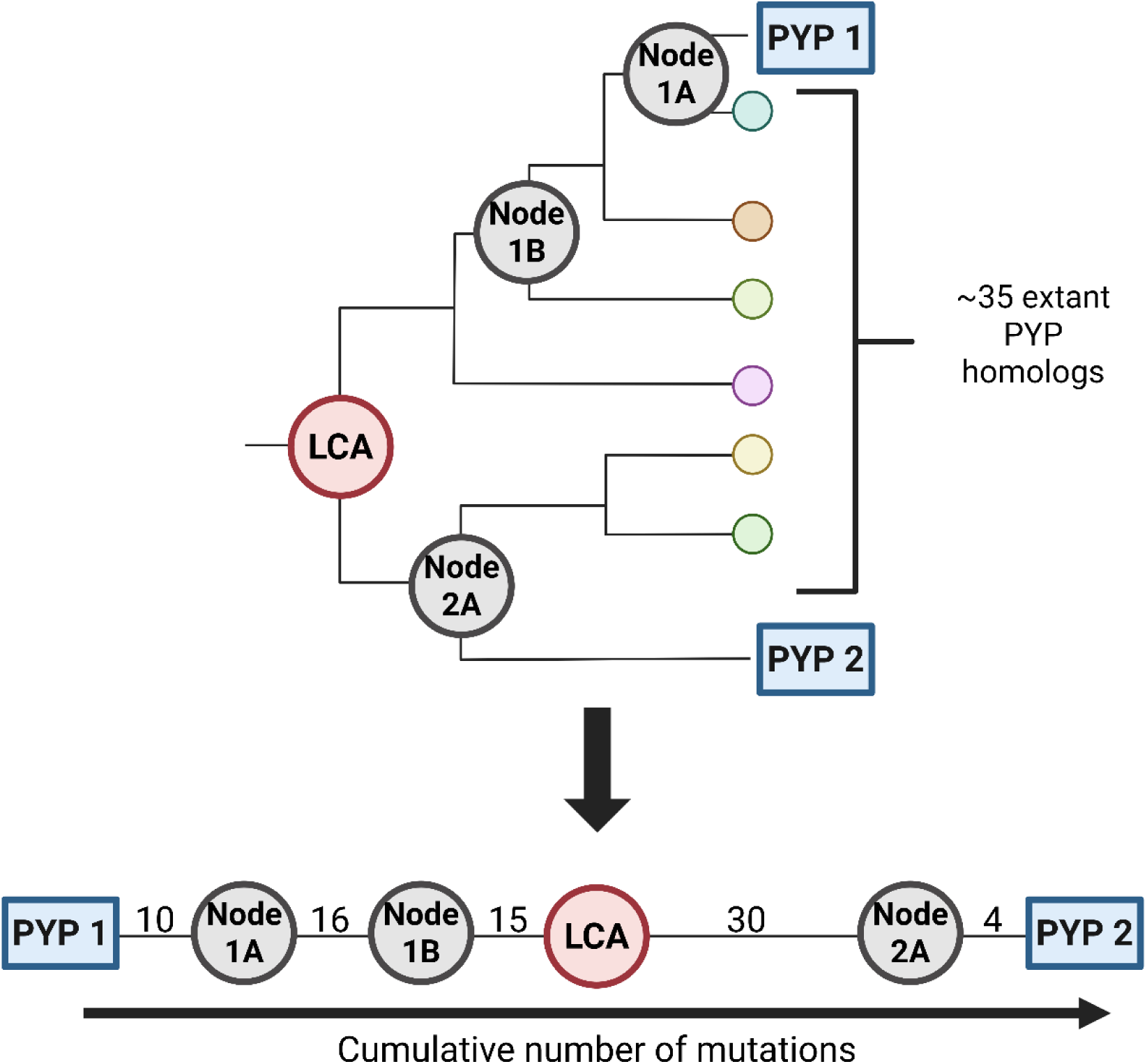
Path through sequence space connecting PYP1 and PYP2 via the four resurrected ancestral PYPs studied here. Top panel: Schematic phylogenetic tree showing the two extant PYPs in blue and their last common ancestor (LCA) in red and the other Nodes 1A, 1B, and 2A in grey. Approximately 35 other extant nodes are present between PYP1 and PYP2. Bottom panel: The information depicted in the top panel provides a path through sequence space connecting PYP1 and PYP2. The number of mutations separating different proteins is indicated.

**Table 1.** Percentage pairwise sequence identity and similarity of ancestral and extant PYPs.

|  | **PYP1** | **Node 1A** | **Node 1B** | **LCA** | **Node 2A** | **PYP2** | **Percent Similarity** |
| --- | --- | --- | --- | --- | --- | --- | --- |
| **PYP1** |  | 96 | 91.2 | 84.8 | 77.6 | 75.2 |  |
| **Node 1A** | 92 |  | 92.8 | 86.4 | 79.2 | 76.8 |  |
| **Node 1B** | 83.2 | 87.2 |  | 92 | 81.6 | 79.2 |  |
| **LCA** | 75.2 | 78.4 | 88 |  | 88 | 85.8 |  |
| **Node 2A** | 61.6 | 61.6 | 66.4 | 76 |  | 97.6 |  |
| **PYP2** | 60 | 58.4 | 63.2 | 72.8 | 96.8 |  |  |
| **Percent Identity** |  |  |  |  |  |  |  |

### Visible absorbance maxima and pB decay rates of resurrected ancestral PYPs

All four ancestral proteins selected for experimental studies were successfully overproduced in *E. coli* and purified with yields similar to that of PYP1. Each ancestor exhibited a strong yellow color and we experimentally quantified their λ_max_ and τ_pB_ (Figure 5, 6 and supplemental Table 1).

A key functional property of PYPs is their visible λ_max_, as they function as blue-light photoreceptors. PYP1 has a λ_max_ of 447 nm while PYP2 displays a slightly blue-shifted λ_max_ at 442 nm. All four resurrected ancestral PYPs exhibited a strong absorbance band with a maximum in the expected spectral region, demonstrating that they fold into a native conformation with intact active site interactions at the *p*CA binding pocket. The resurrected LCA of PYP1 and PYP2 exhibits a λ_max_ of 446 nm. The λ_max_ of the ancestor, Node 2A, in between the LCA and PYP2 is at 442 nm, while the two intermediate ancestors, Node 1A and 1B, in between the LCA and PYP1 both have a λ_max_ of 447nm (Figure 3A and 3B). Thus, PYP1 exhibits the ancestral λ_max,_ while PYP2 evolved to acquire a blue-shifted λ_max_. The extinction coefficients of the *p*CA chromophore in the ancestral PYPs were found to be almost identical the that of the extant protein (Supplemental Figure 2).

**Figure 3.**
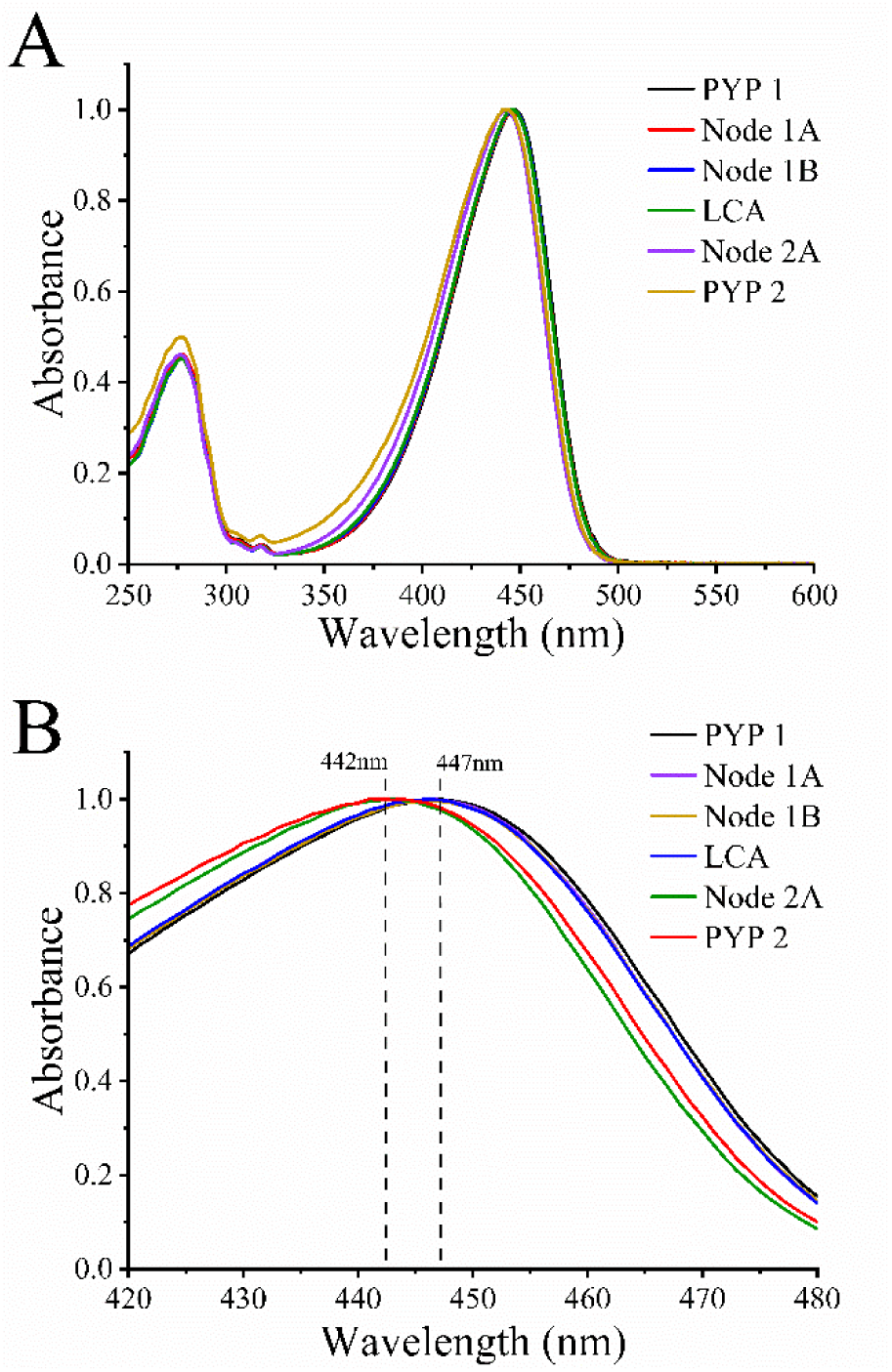
**UV/Vis absorbance spectra of the pG state of ancestral and extant PYPs at room temperature and neutral pH. (**A) Spectra of the PYPs from 250 nm to 600 nm. (B) Same spectra as in A for the spectral region 420 nm to 480 nm to highlight differences in the absorbance maxima (λ_max)_ of this set of PYPs.

To identify residues that are likely to contribute to the blue-shift in the λ_max_ of PYP2, we compared mutations along the path through sequence space from PYP1 to PYP2 (Fig. 2) with the published results on mutations that affect the absorbance spectrum of PYP1. Of the 52 positions at which substitutions occurred along this path through sequence space, only one was at a position known to affect the λ_max_ of PYP1: residue 70 (Fig. 1; supplemental Fig. 1). It is exactly at the transition from the LCA to Node 2B, where a clear blue-shift occurs (from 446 nm to 443 nm), that this residue changes from a Thr to a Gly. For Hhal PYP1 it has been shown that the T50A mutant has λ_max_ of 441 nm ^55^, indicating that the presence of Gly at position 70 in Node 2A and PYP2 results in their somewhat blue-shifted absorbance maxima.

The rate of pB decay is another key functional biochemical property of PYPs. We experimentally measured the value of τ_pB_ for the six PYP variants studied here (Supplemental Table 1). To examine if all these PYPs exhibit the canonical photocycle observed in PYP 1, we measured their UV/Vis light-dark absorbance difference spectra. For PYP1 this difference spectrum has a negative peak near 447 nm caused by the photoconversion of the initial pG state and a positive peak with lower amplitude near 355 nm caused by the formation of the blue-shifted pB photocycle intermediate. With small variations in the relative amplitude of the peak near 355 nm, all PYP variants showed a prominent bleaching of the pG state and the formation of a pB state (Figure 4A and Supplemental Figure 3). These results confirm that that the resurrected ancestors have a functional photocycle and that ancestral reconstruction preserved the essential functional amino acids critical to the protein’s activity.

**Figure 4.**
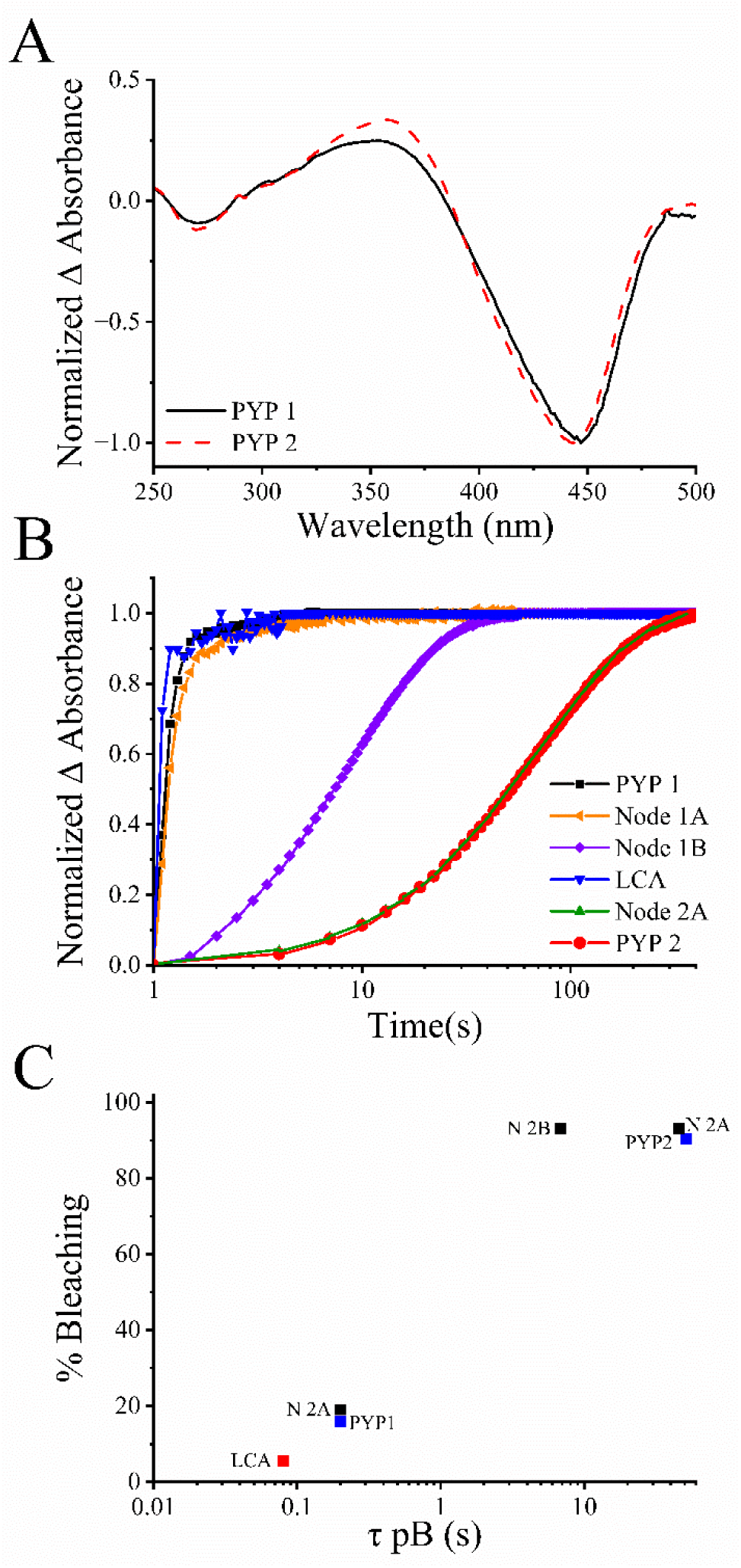
Photocycle characteristics of extant and ancestral PYPs. (A) UV/Vis light-dark difference absorbance spectra of PYP1 and PYP2 reveal spectral changes induced upon continuous illumination. The spectra show the bleaching of the initial pG state near 445 nm (amplitude normalized to 1) and an increase in absorbance near 355 nm, characteristic of the pB photocycle intermediate. (B) The kinetics of pB decay for the indicated PYP variants were measured at 445 nm. To allow direct comparison, the amplitude of all decay traces was normalized to 1. (C) Correlating pB decay rate with % bleaching at 445 nm under continuous illumination (photostationary mixture of pB and pG). Each trace in (B) was fit as a monoexponential decay to derive the lifetime of pB (τ_pB_). The observed correlation between % photostationary bleaching and τ_pB_ validates the pB decay rates depicted in panel (B).

PYP1 exhibits a pB decay rate of 0.5 seconds under the buffer conditions used here, while we measured a τ_pB_ for PYP2 of 52 seconds, very similar to the published value of 77 seconds ^49^. For the resurrected LCA we determined a τ_pB_ of 0.08 seconds, slightly faster than PYP1. Thus, as for λ_max_, PYP1 exhibits the ancestral phenotype and PYP2 diverged to acquire a slower τ_pB_. The intermediate ancestor on the way from the LCA to PYP2 (Node 2A) had a τ_pB_ of 46 seconds, quite close to the value of 52 seconds for PYP2.

To validate the values of τ_pB_ derived from time-resolved UV/Vis absorbance spectroscopy, we employed the correlation between the value of τ_pB_ and the degree of steady-state bleaching of the visible absorbance band of pG under continuous blue-light illumination ^33,72^. The expected trend is that PYP variants with a slower rate of pB decay will exhibit a larger percentage of steady-state bleaching of the pG state. Deviations from this pattern have been reported and indicate the involvement of additional photocycle intermediates ^33,72^. For the six PYP variants studied here we observed that the τ_pB_ correlated well with the percentage photostationary photoconversion of the pG state (Figure 4C), validating the values obtained for τ_pB_.

To examine which substitutions resulted in the change in here τ_pB_ changes from 0.08 seconds in the LCA to 46 seconds in Node 2A, we identified changes in the amino acid sequence of these two ancestral PYPs at positions where substitutions have been shown to alter the τ_pB_ in Hhal PYP1. This analysis yielded six candidate residues: positions 11, 30, 33, 45, 64, 70, and 74 (supplemental Fig 1). For each of these positions an Ala mutagenesis scan showed a modest reduction in the rate of pB decay, providing clear candidates for causing the value of τ_pB_ in PYP2. However, since in PYP2 most of these residues have undergone substitutions to different side chains, more work will be needed to achieve a residue-level explanation.

To examine changes in protein structure during the divergence of PYP1 and 2 from their LCA, we performed AlphaFold2-based structural predictions along the reconstructed evolutionary trajectory to examine if they adopt the typical fold of PYP ^21^. All the predicted structures were of high confidence along their entire sequence. To examine the degree of similarity in this set of predicted structures, pairwise root-mean-square-deviation (RMSD) values were computed with each other and with high resolution crystal structures of PYP ^23,73^, and were found to range from 0.1 to 1.1 Ǻ, centered near 0.55 Ǻ (Figure 5; Table 2). Since RSMD values below 3Ǻ are considered to be indicative of structural conservation ^74^, this set of predicted PYP structures share a very high degree of structural similarity. The RMSD between PYP1 and the PYP homolog from *Rhodospirillum centenum* ^75^ is 1.5 Å (Supplemental Table 2), clearly larger than the values depicted in Figure 3. Furthermore, we analyzed the pairwise RMSD values of 14 high resolution crystal structures of PYP1 ^76^ and found that this set of crystal structures for the same proteins displayed the same range of RMSD values at those for the set of predicted structures for the set of PYPs examined here (Figure 5). No clear correlation was seen between the degree of amino acid sequence (see Table 1) and the RMSD value. We conclude that with respect to their computationally predicted structures, PYP2 and the four ancestral PYPs studied here are all indistinguishable from PYP1 based on their RMSD values.

**Table 2.** Pairwise RMSD values of the three-dimensional structures of PYP1 with PYP2 and the resurrected ancestral proteins studied here. For PYP1 two different experimentally determined crystal structures (1NWZ and 1D7E) were used. To include any effects caused by PYP crystal forms, we examined 1NWZ with space group symmetry p63, and 1D7E with space group symmetry of p65. For PYP2 and the ancestral PYPs predicted structures were used. The structural similarity for each pair of structures was quantified by calculating the root mean square (RMSD) deviation of the positions of all Cα atoms in the protein was computed and depicted in Å.

|  | **PYP1 (1NWZ)** | **Node 1A** | **Node 1B** | **LCA** | **Node 2B** | **PYP2** | **Compared to PDB: 1D7E**<br>**(p65)** |
| --- | --- | --- | --- | --- | --- | --- | --- |
| **PYP1 (1D7E)** | 0.7 | 0.4 | 0.4 | 0.5 | 0.5 | 0.5 |  |
| **Node 1A** | 0.8 |  |  |  |  |  |  |
| **Node 1B** | 1 | 0.3 |  |  |  |  |  |
| **LCA** | 1.1 | 0.3 | 0.2 |  |  |  |  |
| **Node 2A** | 0.9 | 0.6 | 0.5 | 0.4 |  |  |  |
| **PYP2** | 0.9 | 0.5 | 0.4 | 0.4 | 0.1 |  |  |
| **Compared to PDB: 1NWZ (p63)** |  |  |  |  |  |  |  |

To examine the possibility that specific regions in the predicted PYP structures significantly deviate from that of PYP1, we computed residue-resolved root mean square deviations of each Cα atom along the length of PYP (Figure 6 and Supplemental Figure 4). The PYP homolog from *R. centenum* (Rcen) is known to exhibit a somewhat different conformation in the loop connecting β-strand 4 and β-strand 5 near residue 100 ^75^, and this is clearly seen in Figure 6. The alternate placement of this loop was proposed ^75^ to cause the relatively slow pB decay rate of Rcen PYP of 46 seconds ^30^. In comparison, the RMSD profile of the predicted structures of PYP2 and the three ancestral PYPs show only very small signals, except for residues 1 and 2, and residues 17, 18, 19, and 20 with an RMSD value greater than 2. These fluctuations can be understood based on the partially disordered nature of the N-terminal 25 residues of PYP1 ^77^. We conclude that the five calculated structures of PYP homologs and PYP1 are essentially indistinguishable.

### Quantifying the effect of active hydrogen bonding on spectral tuning using FTIR spectroscopy

We used FTIR spectroscopy to experimentally validate the AlphaFold 2 structure predictions. FTIR signals from side chains at protein active sites can be used to obtain local structural insights at high structural resolutions, while measurements of the Amide 1 backbone vibration provide information on the overall secondary structure content of the protein ^78,79^. To enhance the quality of the spectral data, we analyzed the second derivatives of the FTIR spectra measured here ^80^.

Since it has been shown that the strength of the active site hydrogen bond between the *p*CA and Glu46 affects the λ_max_ of PYP ^43,45^, we sought to determine possible changes in the strength of this hydrogen bond in the PYP variants studied here. The frequency of the C=O stretching mode of Glu46 in PYP reports the strength of this hydrogen bond ^81^ and can be directly observed using FTIR spectroscopy ^82,83^. We therefore measured the FTIR spectra of the PYP derivatives studied here (Figure 7). Nodes 1A and 1B share the same λ_max_ as PYP1 and also share the same peak position of the C=O stretching mode of Glu46 at 1726 cm^-1^ (Figure 7C). Moreover, and in line with the expected relationship, Node 2A and PYP2, which both have a λ_max_ of 442 nm, showed a shift in the peak position of the Glu46 C=O stretching mode from 1726 cm^-1^ in PYP1 to 1723 cm^-1^. This shift in the position of the Glu46 C=O stretching mode indicates a strengthening of the Glu46 – *p*CA hydrogen bond, in line with the observed blue-shift in visible λ_max_. The one exception to this pattern is the LCA, which displays a stronger hydrogen bond, evidenced by its Glu46 C=O stretching mode at 1723 cm^-1^ despite having a λ_max_ of 447 nm (Figure 7C). We tentatively attributed the spectral properties of the LCA to subtle structural adjustments that lead to compensatory changes in active-site hydrogen bonding strength ^84^. This observation is intriguing because all of the 15 residues that are immediately adjacent to the pCA chromophore in PYP1 ^7^ are conserved in this LCA, suggesting subtle long-range effects resulting in a structural change at the *p*CA binding pocket. Based on these results we aimed to further examine these suspected structural changes using FTIR spectroscopy in the Amide I region of the spectrum.

**Figure 5.**
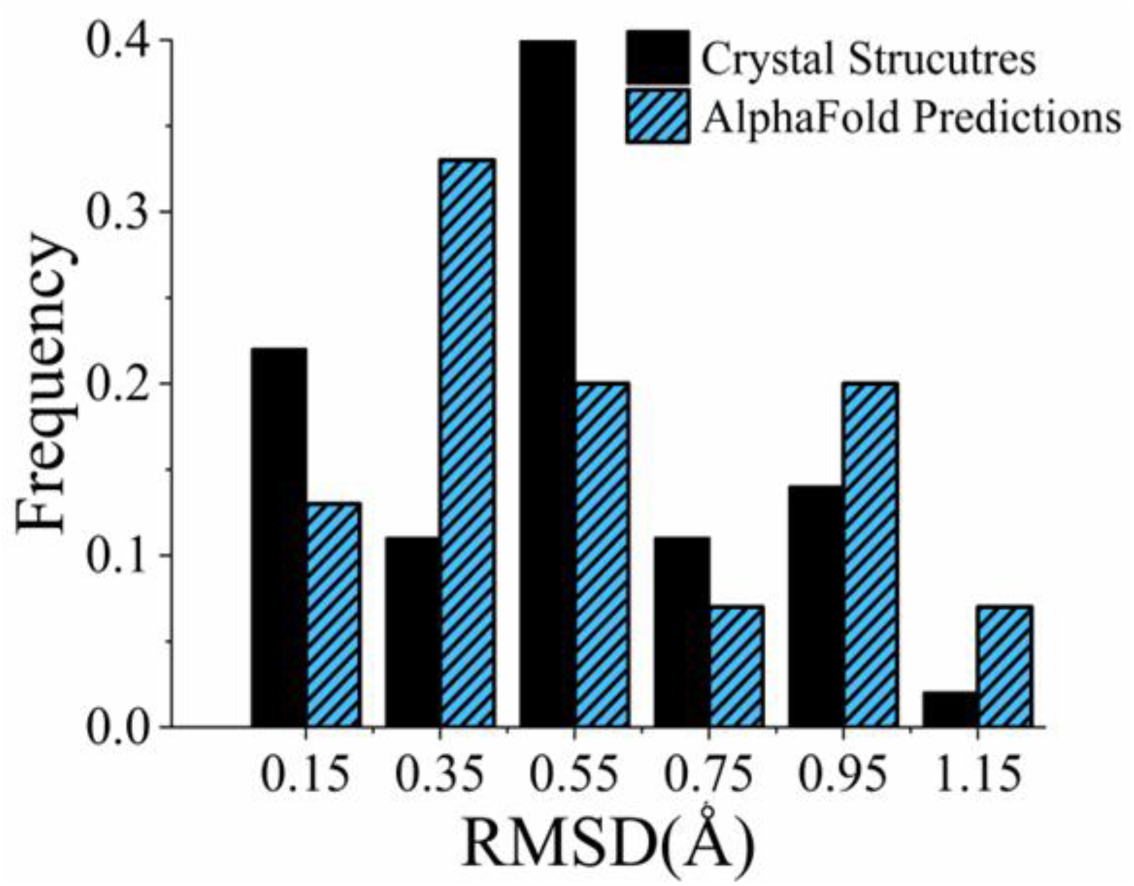
Quantifying changes in the computed three-dimensional structures along the evolutionary pathway resulting in PYP1 and PYP2. Blue bars: The three-dimensional structures of PYP2 and the 4 resurrected ancestral PYPs studied here were computed using AlphaFold2. The pairwise values root mean square (RMSD) deviations of the positions of all Cα atoms (Å) for this set of predicted structures with each other and with PYP1 (PDB ID 1NWZ) was computed. The resulting distribution in pairwise RMSD values is shown. The x-axis depicts the center of each bin. Black bars: To evaluate the distribution of RMSD values of the series of PYP homologs studied here, pairwise RMSD values were computed for a set of 14 independently determined previously reported high resolution crystal structures of the pG state PYP1. The resulting distribution of pairwise RMSD values is depicted.

**Figure 6.**
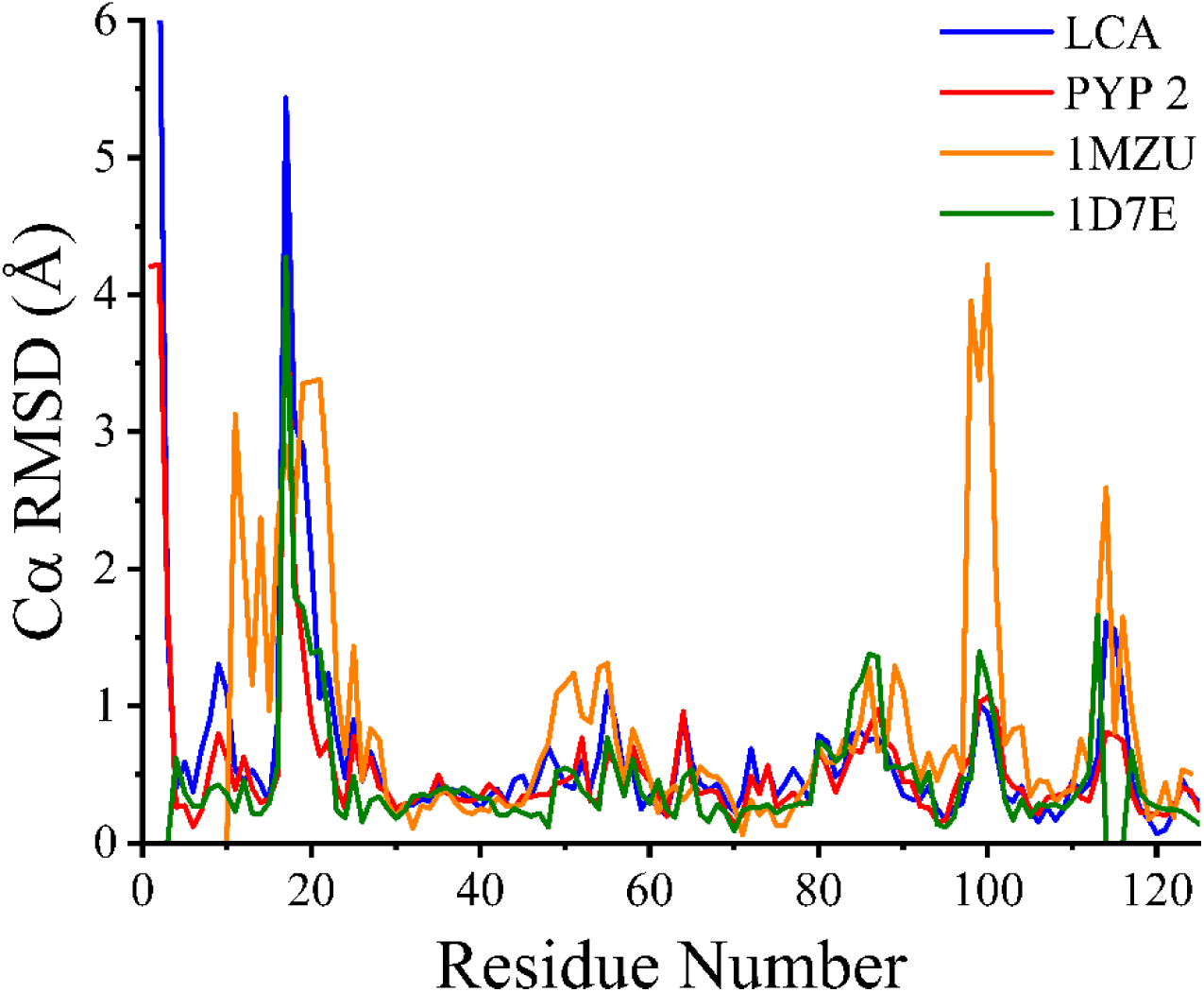
Residue-resolved root mean square fluctuation profiles for PYP2, the last common ancestral PYPs of the pair studied here compared to PYP1. The root mean square deviation (RMSD) of the Cα carbon for each residue in the protein structure is depicted. All four structures were compared with crystal structure 1NWZ (space group p63). The RMSD values for LCA (blue) and PYP2 (red) were obtained using predicted structures. 1MZU is the structure for *Rhodosprillum centenum* PYP (orange) and 1D7E is a structure of PYP1 in space group p65 (green).

**Figure 7.**
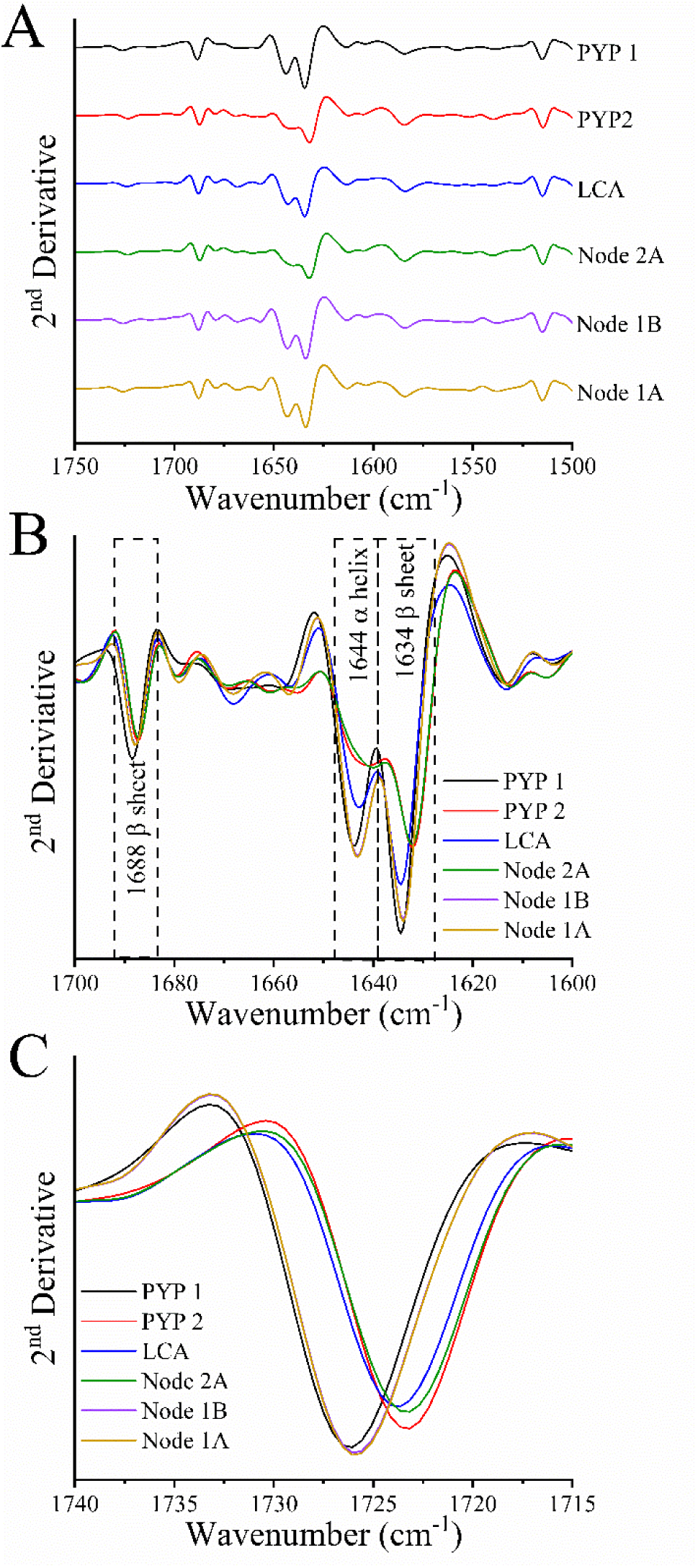
Analysis of the structure of the pG state of ancestral and extant PYPs using second derivative FTIR spectroscopy. (A) Second derivative FTIR spectra for the indicated PYP homologs at room temperature, neutral pH in the spectral region 1750 cm^-1^ to 1500 cm^-1^. (B) Amide I region of the spectra in panel A to examine secondary structure. (C) The spectral region 1740 cm^-1^ to 1715 cm^-1^ contains the characteristic Glu46 C=O stretching mode in these PYPs.

### Detection of notable differences in the secondary structures of ancestral PYPs by FTIR spectroscopy

FTIR spectroscopy can provide important insights into protein secondary structure based on the Amide I vibrational mode, which consists of the C=O stretching modes of all the backbone carbonyl groups of a protein ^63,85^. The peak position of the Amide I mode of α-helices typically is in the range 1648-1657 cm^-1^, while β-sheets generally exhibit two Amide I bands: a stronger band near 1612-1641 cm^-1^ and a weaker band near 1670-1694 cm^-1^.

Here we compare the secondary structures of the six protein variants studied here based on the features of their Amide I absorbance band. Such detailed comparisons of the FTIR spectra of different protein samples often are hampered by technical issues, including shifts in the baseline of the spectra. To overcome this challenge, we used a two-pronged strategy. First, we used an AquaSpec constant path length FTIR sample cell to maximize sample to sample reproducibility ^70^. Second, we probed secondary structure after calculating the second derivative of the measured infrared spectra, which removes most artifacts caused by baseline shifts ^80^. The resulting spectra are shown in Figure 7.

Consistent with the mixed α/β structure of PYP ^11,21,77^, we observed Amide I bands near 1644 cm^-1^ attributable to α-helical structure and bands near 1634 cm^-1^ and 1688 cm^-1^ indicative of β-strands (Figure 7B). However, a striking degree of variation in these Amide I bands was detected. PYP1 exhibits three prominent peaks at 1688 cm^-1^, 1644 cm^-1^, and 1634 cm^-1^. Nodes 1A and 1B share these peaks, consistent with their close sequence similarity to PYP1. In contrast, Node 2A (the LCA) shows a reduction in peak amplitude, suggesting structural divergence in the α-helix. Node 2A and PYP2, the two most closely related by sequence, demonstrate an almost complete loss of the α-helical peak at 1644 cm^-1^ (Figure 7B). It should be noted that the amplitude of peaks in the Amide I region in second derivative spectra is affected by the width of the absorbance band. An increase in the width an absorbance band will result in a lower amplitude in the second derivative spectrum. Such an increase in Amide I bandwidth can be caused by an increase in structural heterogeneity of the protein, causing inhomogeneous broadening of the absorbance band. We tentatively ascribe the loss of amplitude of Amide I bands in the second derivative spectra depicted in Fig. 7B to an increase in structural inhomogeneity in the affected PYP variants. We conclude that the evolutionary divergence of PYP1 and PYP2 involved significant adjustments in the secondary structure of the protein.

FTIR spectroscopy is emerging as a powerful tool for studies on the structural biology of proteins ^16,79,86,87^. It is amenable to time-resolved measurements and can provide both high-resolution information on the hydrogen bonding status and protonation state of side chains and overall secondary structure content of proteins. The results reported here demonstrate the power of FTIR spectroscopy here for detecting structural changes in a set of homologous proteins.

Taken together, the results reported here demonstrate how ASR can provide insights into mechanisms leading to the divergence of members of a protein family, and provide a striking demonstration that substantial adjustments in the protein secondary structure of members of the same protein family can go entirely undetected by structural predictions based on Alphafold2. This conclusion has clear implications for the use of AlphaFold predictions in uncovering the structural basis of the widely occurring phenomenon of functional divergence in protein families. Future experimental work is needed to further study how the experimentally detected divergence in protein structure in the PYP family cause the observed functional changes in the values of τ_pB_ and λ_max_.

## Acknowledgements

This work was supported by the National Institutes of Health (R15GM144898 to A.X. and W.D.H.) and the National Science Foundation (MRI 1338097 to A.X. and W.D.H.; I-Corps 2449014 to W.D.H. and R.D.). Additional support was provided by the Oklahoma State University for the Advanced FT-IR Research Facility from the Office of Vice President for Research, College of Arts and Sciences, and the Department of Physics. We thank Prof. Mostafa Elshahed for valuable intellectual input and stimulating discussions on protein evolution. We thank Dr. Matthew Cabeen for helpful comments on this manuscript. We acknowledge access to FTIR instruments located in the Center of Advance Infrared Biology at Oklahoma State University. We thank Dr. Chelsea Murphy at the High-Performance Computing Center at Oklahoma State University for support with bioinformatics analyses and Dr. Steve Hartson and Janet Rogers at the DNA Sequencing Core Facility at Oklahoma State University.

## Supplemental Materials

### Supplemental methods

#### Homolog Search, Multiple Sequence Alignment and Ancestral Sequence Reconstruction of PYP

The sequence of PYP1 from *H. halophila* (PDB: 1NWZ) was used as the query for an NCBI PSI-BLAST search against the non-redundant database, employing the PAM250 matrix, a word size of 3, and gap costs 14,2, to identify PYP homologs (as of December 2022). Prior to creating the multiple sequence alignment (MSA), the retrieved sequences were filtered to exclude duplicated and synthetic mutant sequences and all sequences were truncated to a length of 145 amino acids. A high-quality multiple sequence alignment was generated using MAFFT version 7.490 ^64^, employing the globalpair method (G-INS-i). Sequence refinement was then performed with CIAlign version 1.0.17 ^65^, which removed insertions present in fewer than 50% of the sequences and eliminated poorly aligned terminal regions. Once complete the MSA was manually inspected to ensure that all homologs contained functionally essential residue Cys69. The resulting high-quality MSA was subsequently used to construct a phylogenetic tree. A maximum likelihood phylogenetic tree of the PYP homologs was generated using IQ-TREE 1.16.12 ^66,67^ with 1000 bootstrap replicates to ensure robustness. As an outgroup, PAS domain protein PDB: 3FG8 from *Rhodococcus jostii* RHA1 was used. Once IQ-TREE has generated a phylogenetic tree of the extant PYPs, this program was used to ancestral sequences of ancestral PYP based on an empirical Bayesian method.

#### ColabFold: Alphafold-based Structure Prediction

ColabFold ^62^ integrates MMseq2 for fast homology searches with AlphaFold2 to improve the efficiency and precision of structure predictions. Protein sequences are entered into the program, which was used with its default configuration. The predicted structures were then examined using PyMol 3.0.4 ^88^. The RMSD was computed using the Dali Server ^68^. Two PDB files were used as an input and the secondary structure alignment including the RMSD value was given as the output. RMSF values were computed using ChimeraX 1.9 **^Error! Cannot open file.^** . The two input PDB structures were aligned and the Cα RSMF value were provided for each residue.

#### Strain Construction for Synthetically Generated Extant and Ancestral PYPs

The genes encoding ancestral and extant PYPs were synthesized by Genscript and cloned into pET28b+ between the NdeI and BamHI restriction sites, which also provides kanamycin resistance for selection. To remove the sequence encoding the N-terminal histidine tag, the plasmid is digested with the restriction enzymes NcoI (R0193T) and BamHI (R0136S) from New England Biolabs (NEB). The insert was amplified via polymerase chain reaction (PCR), followed by Gibson Assembly (E5510S) from NEB to assemble the newly constructed plasmid. The completed plasmid was transformed into NEB® 5-alpha Competent *E. coli* (High Efficiency) cells, and transformants were selected on kanamycin LB plates to select for the assembled plasmid. Plasmid DNA was extracted and sent for sequencing at the DNA Protein Core Facility at Oklahoma State University, where T7 promoter primers were used to confirm the inserts sequence and integrity. Lastly, the plasmid was transformed into *E. coli* strain BL21 (DE3) (C2527H) from NEB for PYP overproduction. This process was repeated for the other synthesized ancestral and extant protein sequences. It is important to note that PYP2 was purified with a 5 amino acid C-terminal tail and with this tail deleted. The λ_max,_ τ_pB_ and extinction coefficient determined by SDS denaturation showed no distinct differences. Because of this result, we proceeded with the PYP2 lacking its 5-residue tail so that all of our constructs were 125 residues long.

#### Growth of E. coli containing various PYP constructs

A Luria-Bertani (LB) agar plate containing 50µg/mL of kanamycin was streaked with our BL21 *E.coli* strain containing the construct from frozen stock and incubated at 37℃ for 16-20 hours or until colonies appeared. To promote aeration 2x 1L of LB broth media with 50µg/mL kanamycin was prepared in a 2.8L flask. The first 1L was inoculated with multiple colonies from the agar plate to create a preculture, which was grown at 37℃ with shaking at 250 rpm for 10-12 hours. After incubation, the OD_600_ of the pre-culture was measured to estimate the cell density. The cells were then pelleted at 3750 rpm (3273 x g) for 20 minutes and resuspended in the second 1L LB flask with 50µg/mL kanamycin as described in ^56^. Once resuspended, the initial OD_600_ of the main culture was measured, and it was then incubated at 37℃ while shaking at 250 rpm. The OD_600_ was continually measured every 45 – 60 minutes until the OD_600_ had increased by 0.8-1.0. Once the desired OD_600_ was reached, the culture was induced with 1mM (final concentration) IPTG and incubated for another 7-8 hours. After incubation, the cells were harvested at 3750 rpm (3273 x g) for 20 minutes. The supernatant was discarded, and the cell pellet was frozen at -80℃ until ready for purification.

#### Protein purification of ancestral and extant PYPs

Cells containing the different ancestral and extant constructed were lysed with 90mL/L of 8M urea, and protease inhibitor (cat: 04693116001) and DNase (cat: 10104159001) both purchased from Millipore Sigma were added to the cell pellet. The mixture was stirred for 1 hour at room temperature until the pellet was fully resuspended. The solution was pelleted at 30,000 x g for 20 minutes at room temperature. The supernatant was collected, and the urea was diluted to a final concentration of 4M with 20mM Tris pH 7.5. Immediately after diluting the solution, 400µl/L of *p*-coumaric acid anhydride was added while rapidly stirring. The reaction was then stirred for 60 minutes at room temperature. The mixture was then dialyzed overnight against 20mM Tris pH 8.5, stored in the dark at 4℃.

The protein solution was applied to a DEAE Sepharose Fast Flow (cat: 17-0709-01 purchased from Cytiva) column pre-equilibrated with 20mM Tris pH8.5. The protein was loaded onto the column and washed with 20Mm Tris pH 8.5. Elution was carried out in steps with 20,40,60, and 80 Mm NaCl in 20mM Tris pH 8.5. Fractions were collected and their absorbance was measured at 280 nm and 446 nm to calculate the purity index (PI) by dividing abs280/abs446. The optimal PI for WT PYP1 is 0.43. The purest fractions were dialyzed overnight in 20mM Tris pH 7.5 at 4℃. The second round of purification was performed using a Q Sepharose Fast Flow (cat:17-0510-01 from Cytiva) column pre-equilibrated with 20 mM Tris pH 7.5. The protein was loaded onto the column, washed and eluted with 20, 40, 60, and 80mM NaCl in 20mM Tris pH 7.5. Fractions were collected and the purest ones were dialyzed against 20mM Bis-Tris pH 6.0 overnight at 4℃. For the final purification step, the Q Sepharose High Performance (cat: 17-1014-01 from Cytiva) column was pre-equilibrated with 20Mm Bis-Tris pH 6.0. The protein was loaded, washed, and eluted with 20, 40, 60, 80Mm NaCl in 20 Mm Bis-Tris pH 6.0. Absorbance spectra of the fractions were measured, and the pure protein was dialyzed against 20Mm Tris pH 7.5 overnight at 4℃. The protein was aliquoted and stored at - 80℃.

#### Photocycle measurements and absorbance spectra

The PYP sample was diluted to an OD_600_ range of 0.3-0.8 in 20mM Tris pH 7.5. Absorbance maximum and photocycle measurements were conducted using a HP-8453 (Hewlett-Packard) diode array spectrophotometer. The sample was placed in the spectrophotometer’s sample holder, which had been previously blanked with 20mM Tris buffer at pH 7.5. For absorbance spectra, the spectrophotometer was operated in standard mode. To measure the photocycle, the spectrophotometer was switched to kinetic mode, with an integration time set to 0.1 seconds. The kinetic measurement was conducted over 5 minutes: after approximately 10 seconds the sample was illuminated with bright light for 5 seconds. Following illumination, the sample was allowed to recover in the dark.

#### Determination the extinction coefficient of PYP using SDS denaturation

The estimation of the extinction coefficient of PYP was determined by measuring the absorbance spectrum before and after denaturation with 2% SDS. The protocol published in was used to determine these values.

#### Protein sample preparation for FTIR measurements

A 3mg aliquot of a pure PYP sample was thawed on ice for preparation. The protein samples were concentrated using a 0.5ml Amicon centrifugal filter with a molecular weight of 10,000, obtained from Millipore Sigma (cat: UFC501096). After prewashing the filter, 0.5 ml of the protein solution was added to it and centrifuged for 5 minutes at 9,000 x g, followed by 7 minutes at 11,000 x g. if the initial sample volume exceeded 0.5 ml, additional sample was added as the volume reduced during centrifugation, until all 3 mg were added. The final protein volume was maintained between 30-45 µl to prevent filter damage due to increased pressure.

The buffer was then exchanged to a solution containing 10mM K_2_HPO_4_ and 10mM KH_2_PO_4_ pH 7.5. For this step, 450 µl of the buffer was added to the ∼30µl concentrated protein sample in a fresh centrifugal filter, and the protein was washed three times with this buffer. The filtrate’s pH was confirmed using an Orion™ Double Junction pH Electrode from Thermo Fisher Scientific (cat: 9110DJWP) to ensure it matched the buffer’s pH. The protein was further concentrated to approximately 5 mM PYP, collected by inverting the filter unit, and centrifuged into a collection tube.

The total sample volume was measured with a Hamilton syringe, frozen with liquid nitrogen and freeze dried overnight. The final PYP concentration for FTIR experiments was adjusted to 10Mm by rehydrating the freeze-dried sample with half the original volume of pure D_2_O.

#### FTIR data collection

Single-beam FTIR spectra of PYP variants in the dark (without photoexcitation) were collected at 2 cm^-1^ resolution from 950 to 4000 cm^-1^ using a Bruker Vertex 80V FTIR spectrometer with a KBr beamsplitter and a liquid nitrogen cooled photovoltaic mercury cadmium telluride (MCT) infrared detector. An AquaSpec (Bruker) FTIR sample cell with fixed sample thickness (∼7.0 μm) was used in all data collection to enhance reproducibility of the FTIR measurements. Two tele-arms of the AquaSpec cell were purged with dry nitrogen gas to remove water vapor prior to and during data collection, to reduce infrared absorption of water vapor. The remaining optical path was in vacuum (6 hPa) that eliminates contributions from water vapor. Each single-beam spectrum of a protein sample was averaged over 256 scans while the single-beam background spectrum of a reference sample was averaged over 1024 scans. The rapid-scan function was used to accelerate the data collection at a mirror velocity of 2.5 cm per second (80 kHz) (check), approximately 1 minute per single beam spectrum of 256 scans. The sample temperature was stabilized at 20.0°C using a thermostatted water bath (Huber, Germany). The noise level in the FTIR data reported here is approximately 0.01 milli-OD.

#### FTIR data analysis

FTIR data processing was performed by first averaging spectra, performing a baseline shift (using a single offset value), then normalizing the amplitude of the resulting spectrum, and finally calculating its second derivative, averaging, and normalization. The Bruker OPUS software was used for these steps. The absorption spectrum of a protein sample, *A*(ν), was calculated using *A*(*ν*) = −*log*_10_[*S*(*ν*)/*R*(*ν*)] where S(ν) denotes a single beam spectrum of the sample, while R(ν) stands for the corresponding single beam spectrum of the reference. Next, averaging four repeatedly measured FTIR absorption spectra yielded not only the averaged absorption spectrum, but also the standard deviation (noise level) of the averaged infrared absorption spectrum. The second derivative of the averaged spectrum was Savitsky-Golay algorithm over 17 data points. To compare FTIR spectra of two protein samples, one infrared spectrum was normalized to the infrared spectrum of another sample by introducing and optimizing a scaling factor in dynamic analysis in OPUS.

---

**Supplemental Figure 1.**
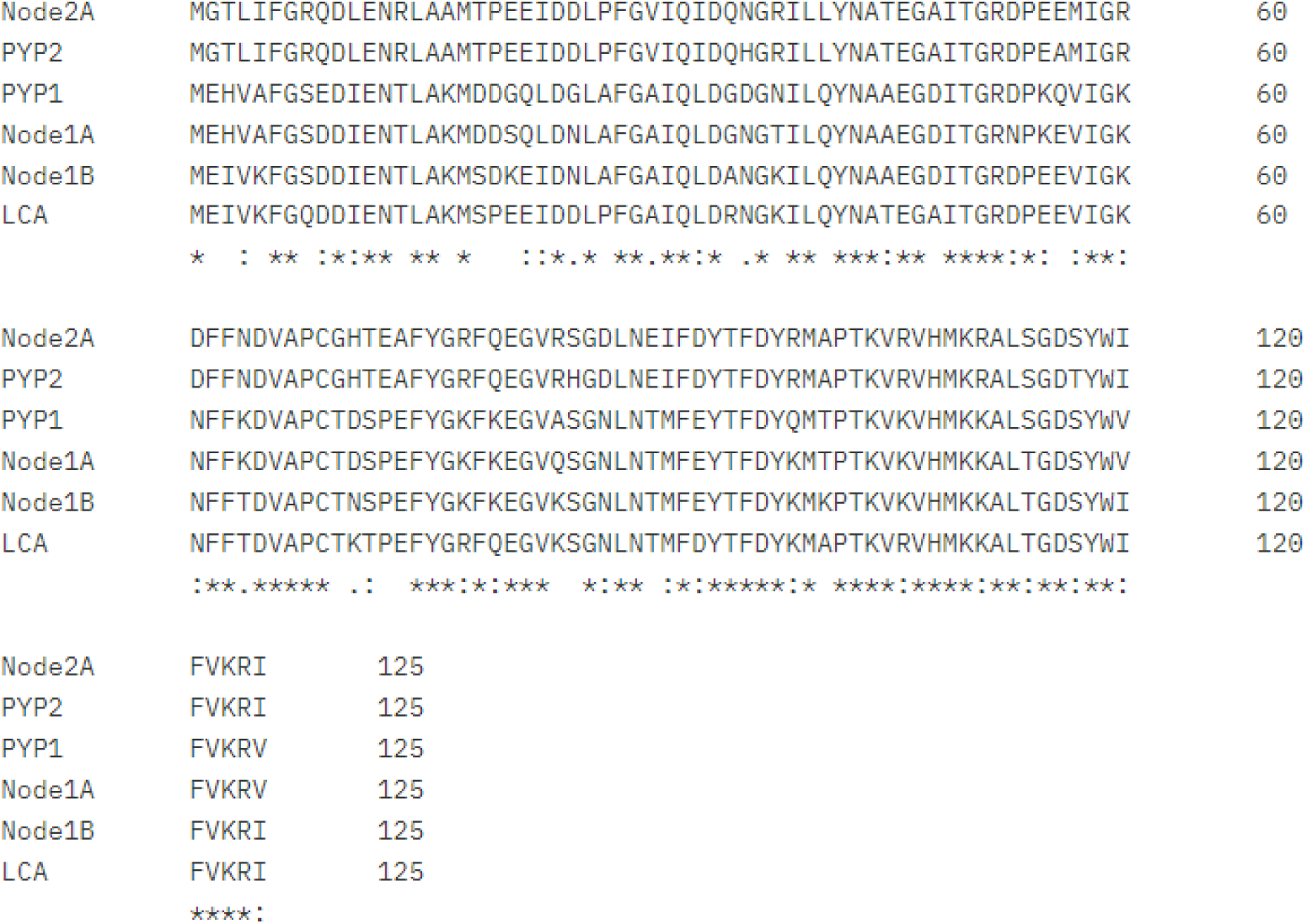
**Sequence alignment of extant PYP and the ancestral nodes that were reconstructed.**

**Supplemental Figure 2.**
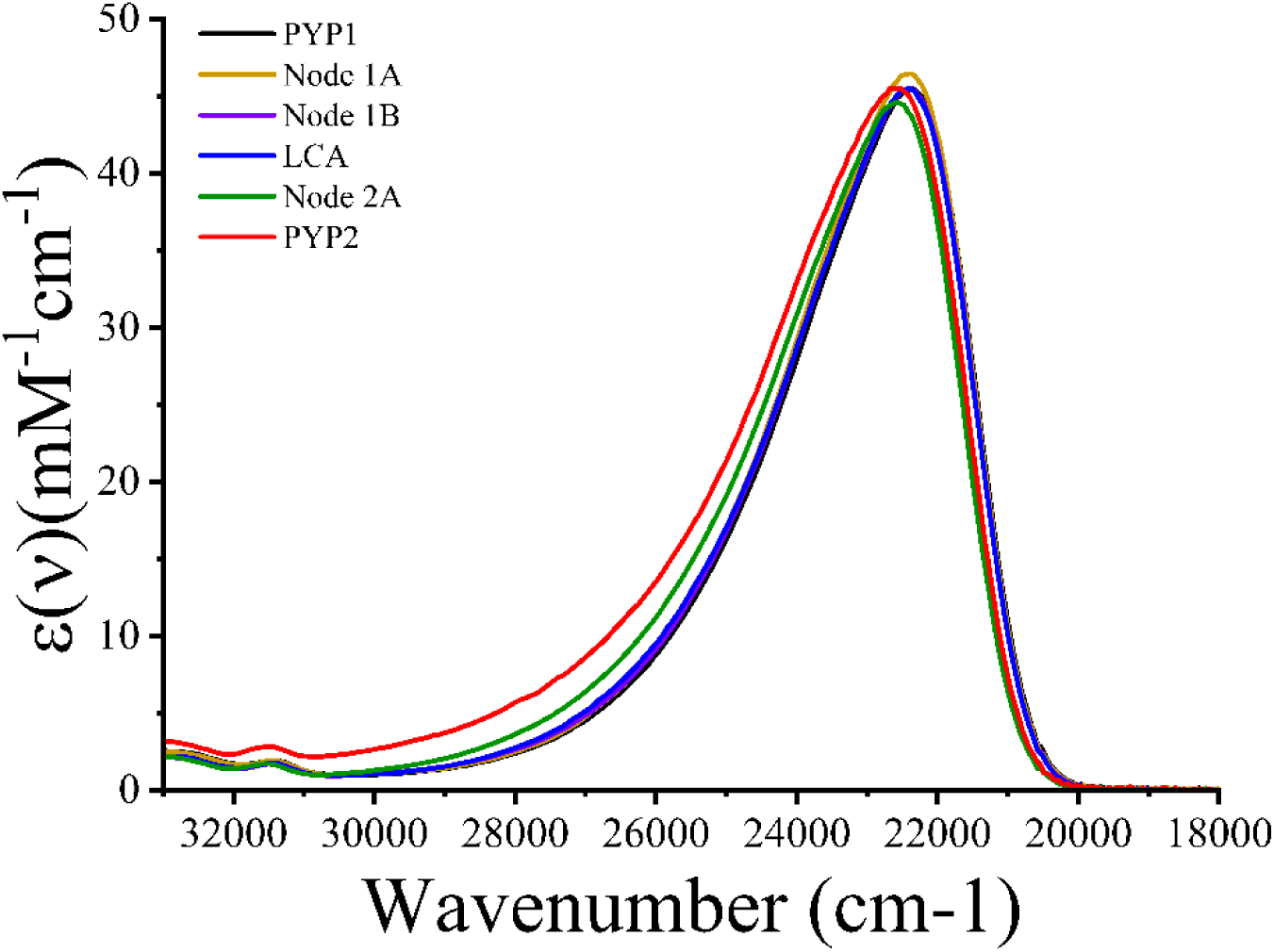
U**V/Vis absorbance spectra of the extant and resurrected ancestral PYPs studied here.**

**Supplemental Figure 3.**
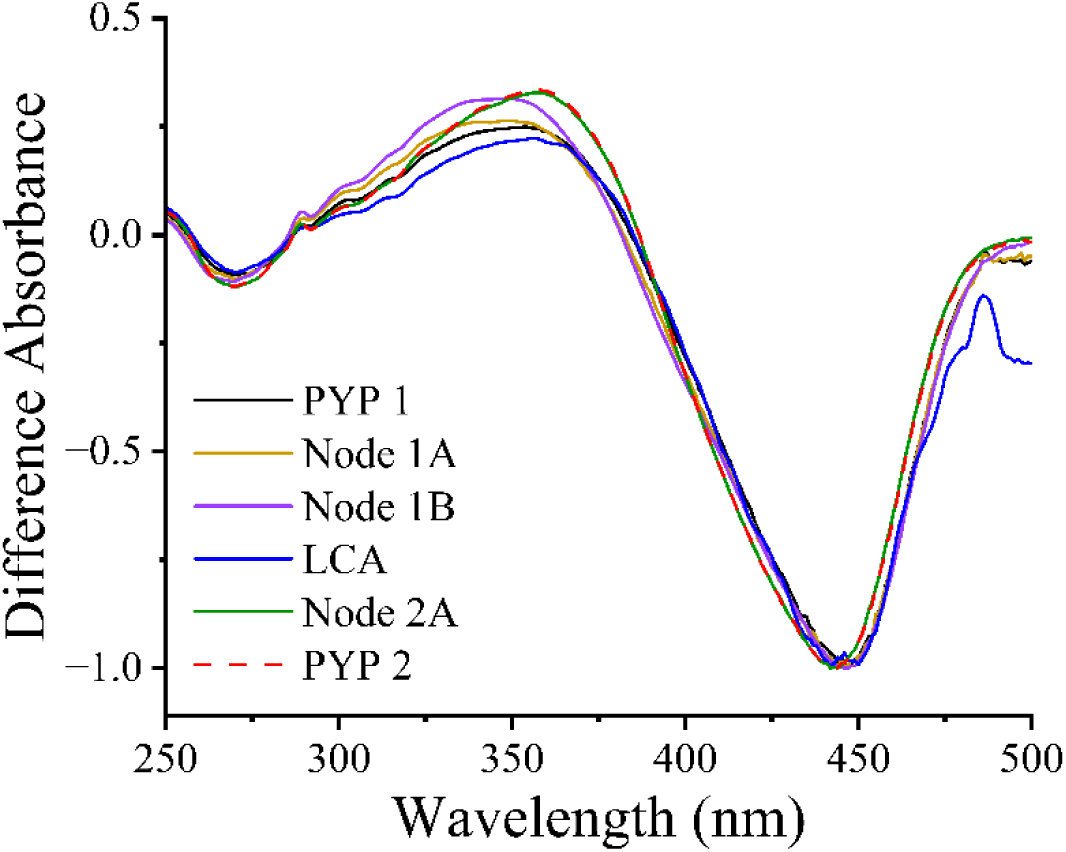
U**V/Vis light – dark absorbance difference spectra of extant and resurrected ancestral PYPs.** The amplitudes of the spectra were normalized to -1 at λ_max_. The signal to noise ratio for the difference spectrum was lower for PYPs with a faster pB decay rate (particularly LCA) because of the small amplitude of the measured difference spectrum.

**Supplemental Figure 4.**
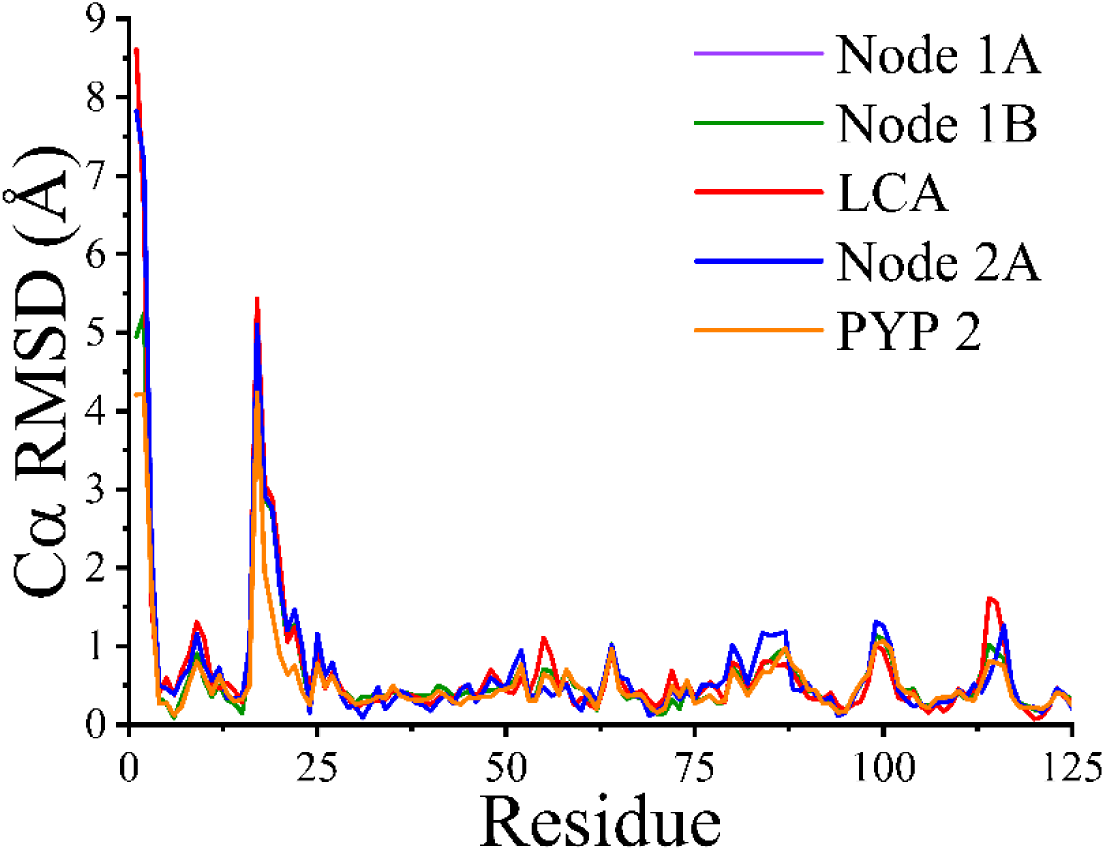
Residue-resolved root mean square fluctuation profiles for PYP2, and the ancestral PYPs of the pair studied here compared to PYP1. The root mean square deviation (RMSD) of the Cα carbon for each residue in the protein structure is depicted. All four structures were compared with crystal structure 1NWZ (space group p63). The RMSD values for Node 1A (purple), Node 1B (green), LCA (red), Node 2A (blue) and PYP2 (orange) were obtained using predicted structures.

**Supplemental Table 1.**
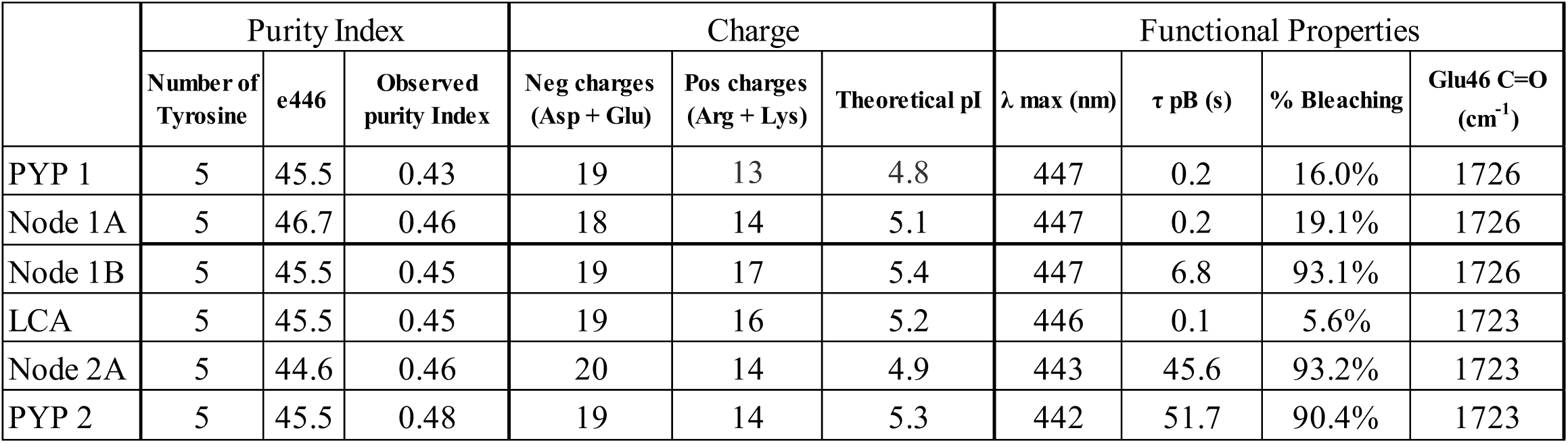
Values of the biochemical data for the PYP variants studied here. The indicated values show the number of tyrosine residues in each PYP, the value of the extinction coefficient at λ_max_ (e446) in mM^-1^cm^-1^ determined using the SDS denaturation method, the highest purity index (ratio of absorbance for the absorbance peaks near 280 nm and near 445 nm) observed upon purification of the samples, the number of positively and negatively charged residues, the computed isoelectric point (pI), the position of the visible absorbance λ_max_ (nm), the lifetime of the pB photocycle intermediate τ_pB_ (seconds), the degree of bleaching of the visible absorbance maximum under continuous illumination, and the position of the C=O stretching mode of Glu46 (cm^-1^) as determined from second derivative FTIR absorbance spectroscopy. Values for PYP 2 were measured for the protein lacking its 5-residue N-terminal extension.

**Supplemental Table 2.**
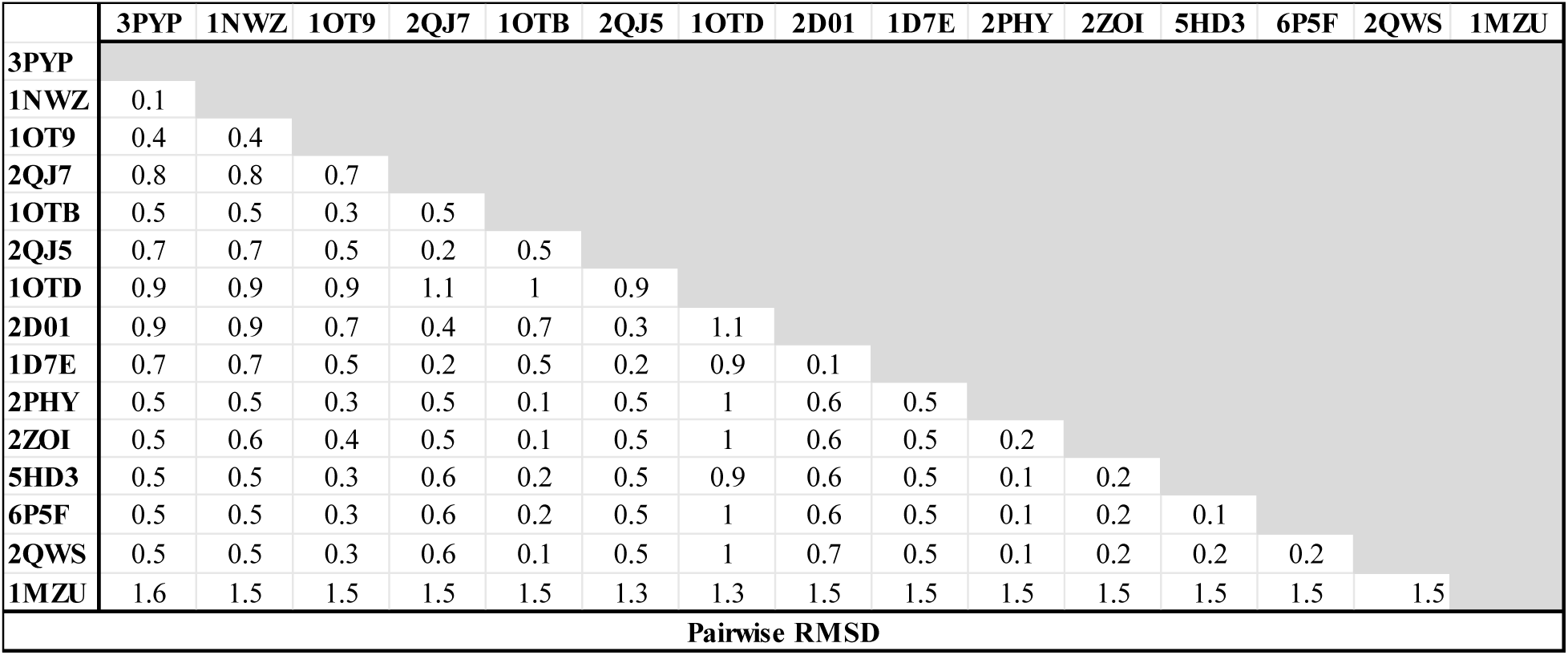
Pairwise RMSD values for a set of experimentally determined crystal structures of the pG state of PYP1. The crystal structure with PDB ID 1MZU is the PYP domain from *Rhodospirillum centenum*.

